# Transcranial alternating current stimulation modulates auditory temporal resolution in elderly people

**DOI:** 10.1101/241471

**Authors:** Alina Baltus, Christoph Siegfried Herrmann

## Abstract

Recent research provides evidence for a functional role of brain oscillations for perception. For example, auditory temporal resolution seems to be linked to individual gamma frequency of auditory cortex. Individual gamma frequency not only correlates with performance in between-channel gap detection tasks but can be modulated via auditory transcranial alternating current stimulation. Modulation of individual gamma frequency is accompanied by an improvement in gap detection performance. Aging changes electrophysiological frequency components and sensory processing mechanisms. Therefore, we conducted a study to investigate the link between individual gamma frequency and gap detection performance in elderly people using auditory transcranial alternating current stimulation. In a within-subject design, nine participants were electrically stimulated with two individualized transcranial alternating current stimulation frequencies: 3 Hz above their individual gamma frequency (experimental condition) and 4 Hz below their individual gamma frequency (control condition) while they were performing a between-channel gap detection task. As expected, individual gamma frequencies correlated significantly with gap detection performance at baseline and in the experimental condition, transcranial alternating current stimulation modulated gap detection performance. In the control condition, stimulation did not modulate gap detection performance. In addition, in elderly, the effect of transcranial alternating current stimulation on auditory temporal resolution seems to be dependent on endogenous frequencies in auditory cortex: elderlies with slower individual gamma frequencies and lower auditory temporal resolution profit from auditory transcranial alternating current stimulation and show increased gap detection performance during stimulation. Our results strongly suggest individualized transcranial alternating current stimulation protocols for successful modulation of performance.

## Introduction

A general decline of sensory processing speed is observed in elderly people (Cerella, 1990; Cerella & Hale, 1994; Li *et al*., 2001; Humes *et al*., 2012). For example, in auditory processing, speech intelligibility drops significantly faster in older adults than in younger adults if speech rate increases (Vaughan & Letowski, 1997). Increasing the rate of speech affects the stimulus in two ways: First, the frequency of contextual information is higher, meaning more bits of information per time have to be processed. Second, temporal acoustic cues are shortened and acoustic information is distorted (Gordon-Salant & Fitzgibbons, 2001). Especially the second aspect seems to be relevant for the decrease in speech intelligibility observed in older adults. Speech comprehension difficulties in older adults are assumed to be the result of a decline in processability of acoustic distortions in accelerated speech rather than a general slowing down in cognitive processing speed (Schneider *et al*., 2005). Aging is assumed to influence the processability of specific temporal cues which are provided by characteristics of consonants, vowels and short intermissions (gaps) in speech (Gordon-Salant & Fitzgibbons, 2001; Gordon-Salant *et al*., 2006). The detection of short temporal intermissions (gaps) in the envelope of speech is pivotal for a successful segmentation of speech. Gaps, as salient points in speech, are assumed to be used as ‘edges’ to convert speech into meaningful, linguistic chunks of information to facilitate further processing (Giraud & Poeppel, 2012). This idea is supported by a significantly diminished speech intelligibility if gaps in speech are artificially shortened by flattening the envelopes of transitions between syllables (Doelling *et al*., 2014).

In general, temporal acuity of auditory processing mechanism seems to be an important aspect for successful speech processing (Phillips, 1999). This leads to the idea that higher temporal acuity and better performance in understanding normal and distorted speech are linkable. For a broader understanding of temporal processes and acuity in auditory perception, gap detection thresholds determined in gap detection tasks are used as behavioral measures of auditory temporal acuity and temporal resolution of the auditory system (Phillips *et al*., 1997). In such tasks, gap detection performance not only varies considerably between younger and older adults (Snell, 1997; Barsz *et al*., 2002) but also between individuals within the same age group (Baltus & Herrmann, 2015; Baltus *et al*., 2017).

A growing body of evidence suggests an interaction between processing mechanisms that are related to processing of short, near-threshold stimuli and frequency characteristics of neuronal oscillations in the brain (Engel *et al*., 2001; Busch *et al*., 2009; Neuling *et al*., 2012a). It is assumed that frequencies of neuronal oscillations in specific brain areas determine temporal resolution of associated perceptual systems. In vision as well as in audition, it has been observed on individual basis that higher frequencies of neuronal oscillations in respective areas led to finer temporal resolution of perception (Baltus & Herrmann, 2015; Samaha & Postle, 2015). Evidence from different perceptual systems suggest a dependency of processing of near-threshold stimuli on distinct phases of neuronal oscillations (Busch *et al*., 2009; Neuling *et al*., 2012a; Riecke, 2016; ten Oever *et al*., 2016). During one cycle of a neuronal oscillation, the respective neuronal ensemble switches once between a more excitable and a less excitable phase causing discrete processing windows which are assumed to shape perception (Engel *et al*., 2001; VanRullen & Koch, 2003; Fingelkurts & Fingelkurts, 2006). In this scenario, the frequency of the neuronal oscillation determines processing rate of sensory events with faster oscillations leading to higher temporal resolution of perception.

A large body of evidence supports the idea of a decisive role of ongoing gamma band oscillations (30 – 70 Hz) in auditory cortex areas for auditory processing (Brosch *et al*., 2002; Lakatos *et al*., 2005; Traub *et al*., 2005; Schadow *et al*., 2007; Cervenka *et al*., 2013). Recent research provides evidence for the assumption that not only different frequency bands can be associated with different perceptual processes but also frequency differences within the gamma band are physiologically relevant on an individual basis (Zaehle *et al*., 2010; Baltus & Herrmann, 2015, 2016; Baltus *et al*., 2017). Unlike in vision, in audition the determination of frequency of relevant neuronal oscillations cannot be estimated from resting state activity but must be derived from indirect measures (Lampl, I., Yarom, 1997). An estimate of the frequency of relevant neuronal oscillations in auditory cortex can be identified using resonance behavior of the auditory cortex elicited by rhythmic stimulation. In an auditory steady state response (ASSR) paradigm, sensory stimulation with amplitude modulated tones (modulation frequencies in the gamma range) leads to an electrophysiological ASSR that follows the stimulation frequency (Picton *et al*., 2003). Bi-hemispheric cortical sources lead to most pronounced ASSR amplitudes at central electrode positions (Herdman *et al*., 2002; Popescu *et al*., 2008). ASSR amplitudes not only vary depending on sensor position but show frequency specificity regarding the modulation frequency with highest amplitudes between 40 and 60 Hz (Zaehle *et al*., 2010). The modulation frequency that elicits the largest ASSR amplitude identifies the characteristic frequency of the neuronal oscillations in the auditory cortex: the individual gamma frequency (IGF). Comparison of IGFs between individuals and within individuals reveals a high inter-subject variability but high intra-subject test-retest reliability (Fründ *et al*., 2007; Baltus & Herrmann, 2015). Recent research provides evidence for a physiological relevance of the IGF for auditory processing in terms of an established correlation between individually determined IGFs and individually determined gap detection thresholds in a between-channel gap detection task (Baltus & Herrmann, 2015). The purely correlational link was supported by a second study where an application of frequency-specific auditory transcranial alternating current stimulation (tACS) during execution of the same between-channel gap detection task led to a frequency-specific modulation of gap detection thresholds on an individual level (Baltus *et al*., 2017). Applying tACS is assumed to modulate ongoing neuronal oscillations in targeted areas by shifting the frequency of the oscillation away from the eigenfrequency (Thut *et al*., 2011; Herrmann *et al*., 2013). Finite element method simulation of tACS current flow allow to investigate current strength and current directions depending on tACS electrode locations (Neuling *et al*., 2012b; Wagner *et al*., 2015; Wagner *et al*., 2016a). A tACS electrode setup specifically suited to stimulate targets in auditory cortices is introduced in Baltus *et al*. (2017). In addition to the electrode locations, the proportions of frequency of the endogenous neuronal oscillation, tACS frequency, and stimulation strength are also crucial if the modulation of the frequency of endogenous neural oscillations is aspired (Notbohm *et al*., 2016). The closer the stimulation frequency is to the endogenous frequency, the weaker the stimulation intensity needs to be in order to modulate the frequency of the endogenous neuronal oscillations (i.e., concept of the Arnold tongue, Pikovsky & Rosenblum, 2003).

Since elderly people seem to be affected in particular by a slowing of processing speed and a decline in temporal acuity, this group of people would probably benefit strongly from a tACS-induced improvement of temporal acuity. Therefore, the current study aimed to provide evidence for a beneficial effect of frequency-specific auditory tACS on auditory temporal resolution in older adults. The second aspect of the current study was to further investigate the link between gap-detection performance and oscillatory activity in the auditory cortex with the focus on older adults. To achieve the purposes of the current study, gap detection thresholds as well as individual gamma frequencies (IGFs) were examined and correlated. IGFs where then used to determine tACS frequencies. Based on the idea that auditory tACS can increase IGF and higher IGFs are correlated with better gap detection performance, we hypothesized an increase in gap detection performance of elderly participants during the experimental condition (A-tACS at frequencies *above* IGF) and no change of the same participants during the control condition (B-tACS at frequencies *below* IGF). In the experimental condition, we were indeed able to modulate gap detection thresholds. The direction of modulation correlated significantly with the IGF, indicating a dependency of the effect of auditory tACS on auditory temporal resolution on endogenous frequencies in auditory cortex in elderly people. In the control condition, as hypothesized, gap detection thresholds were unaffected by auditory tACS.

## Methods

### Participants

In the experiment, twelve participants (six females, mean age: 72 ± 3.1 years) were measured and the data of nine subjects (five females, mean age: 71 ± 3.5 years) were analyzed. Three subjects had to be excluded (two due to technical problems during recording and one participant reported hearing problems in daily situations). All participants took part in the experimental condition and in the control condition. The order of conditions was counterbalanced across all participants. The order of conditions among female and male participants was also randomized. Ten of twelve participants were recruited from the data base of the "Hörzentrum Oldenburg” and pre-screened with pure tone audiometry (PTA) using air conduction (age of PTAs: 3.7 ± 1.8 years). Hearing thresholds were obtained for frequencies from 125 Hz to 4 kHz. All participants were rated by the “Hörzentrum Oldenburg” as normal hearing (see average PTA for details, Fig. 1). Written informed consent was obtained from each participant and the study was approved by the local ethics committee (“Commission for Research Impact Assessment and Ethics”) of the University of Oldenburg, Germany. All procedures were performed in accordance with the Declaration of Helsinki. A short questionnaire was used to obtain information on psychological or neurological disorders, head injuries in the recent past, the occurrence of epilepsy in the personal or family history, and hearing problems in daily situations. No participant reported any of these exclusion criteria, despite the one participant who was excluded from the analysis. All participants were naïve regarding the aim of the study and were untrained in the GD task.

**Figure 1:**
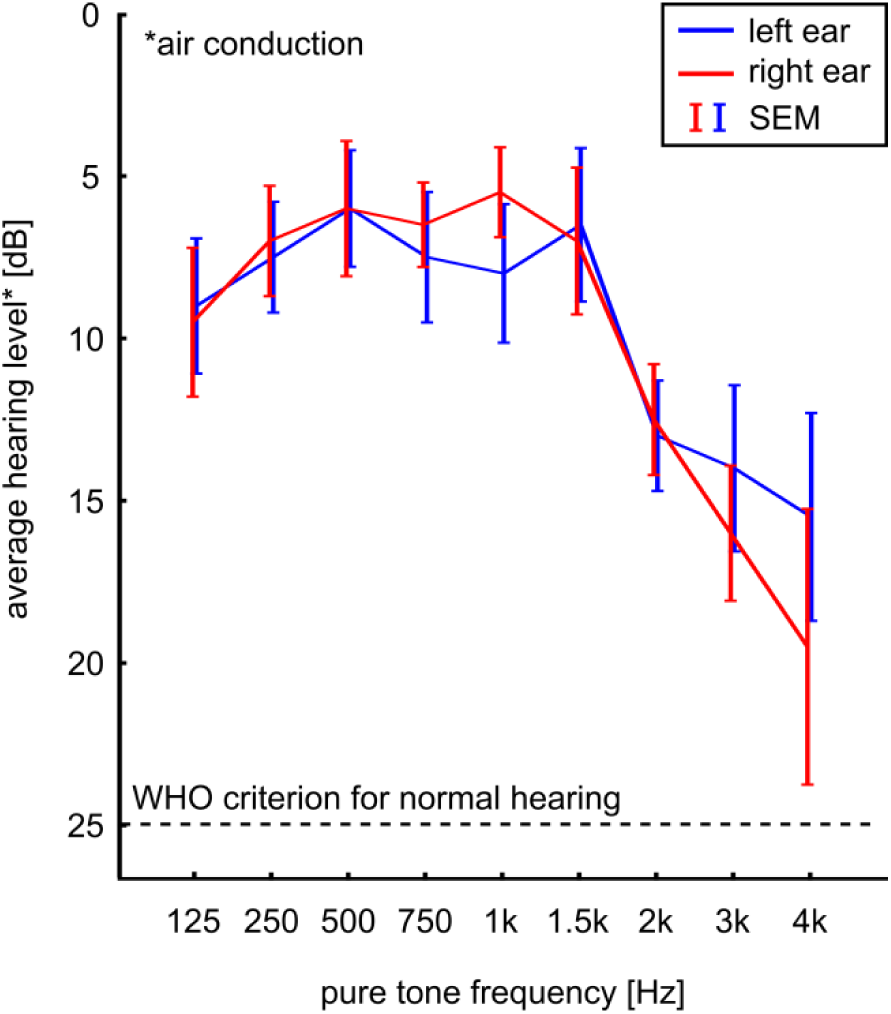
Average pure tone audiograms of nine subjects (age range: 64 to 75 years). Error bars indicate standard error of the mean. Pure tone audiograms were conducted using headphones (air conduction) instead of bone conduction. All subjects were above 25 dB hearing level, the World Health Organization (WHO) criterion for normal hearing.

### Electroencephalography

The experiment was carried out in an electrically shielded room to minimize the influence of 50-Hz line noise. A battery driven light ensured a dimly lit room. EEG was recorded with 3 sintered Ag/AgCl-electrodes (Fz, Cz and Pz) attached to an elastic cap (Easycap GmbH, Herrsching-Breitbrunn, Germany), a nose-tip reference and a ground electrode at AFz (see Fig. 1B1). Impedances were kept below 10 kQ for each electrode. The EEG signal was digitized with the BrainAmp system and recorded with the BrainVision recorder (BrainProducts, Munich, Germany). Sampling rate for digitization was 5000 Hz and the resolution for the amplification was 0.1 μV.

### Experimental setup

The experiment was carried out on two days (see Fig. 2A for graphical depiction of the experimental procedure). Hearing thresholds of all participants were estimated with an adapted version of the self-adjustable adaptation procedure from Carhart and Jerger (Carhart & Jerger, 1959) at day one before the actual experiment started. A white noise burst was used instead of pure tones to determine a rough, frequency unspecific absolute hearing threshold for all participants. Auditory stimuli were then presented in relation to the individual hearing threshold. For validation of this procedure, hearing thresholds for all participants were correlated with the mean individual hearing thresholds obtained with the PTA at the “Hörzentrum Oldenburg”.

**Figure 2:**
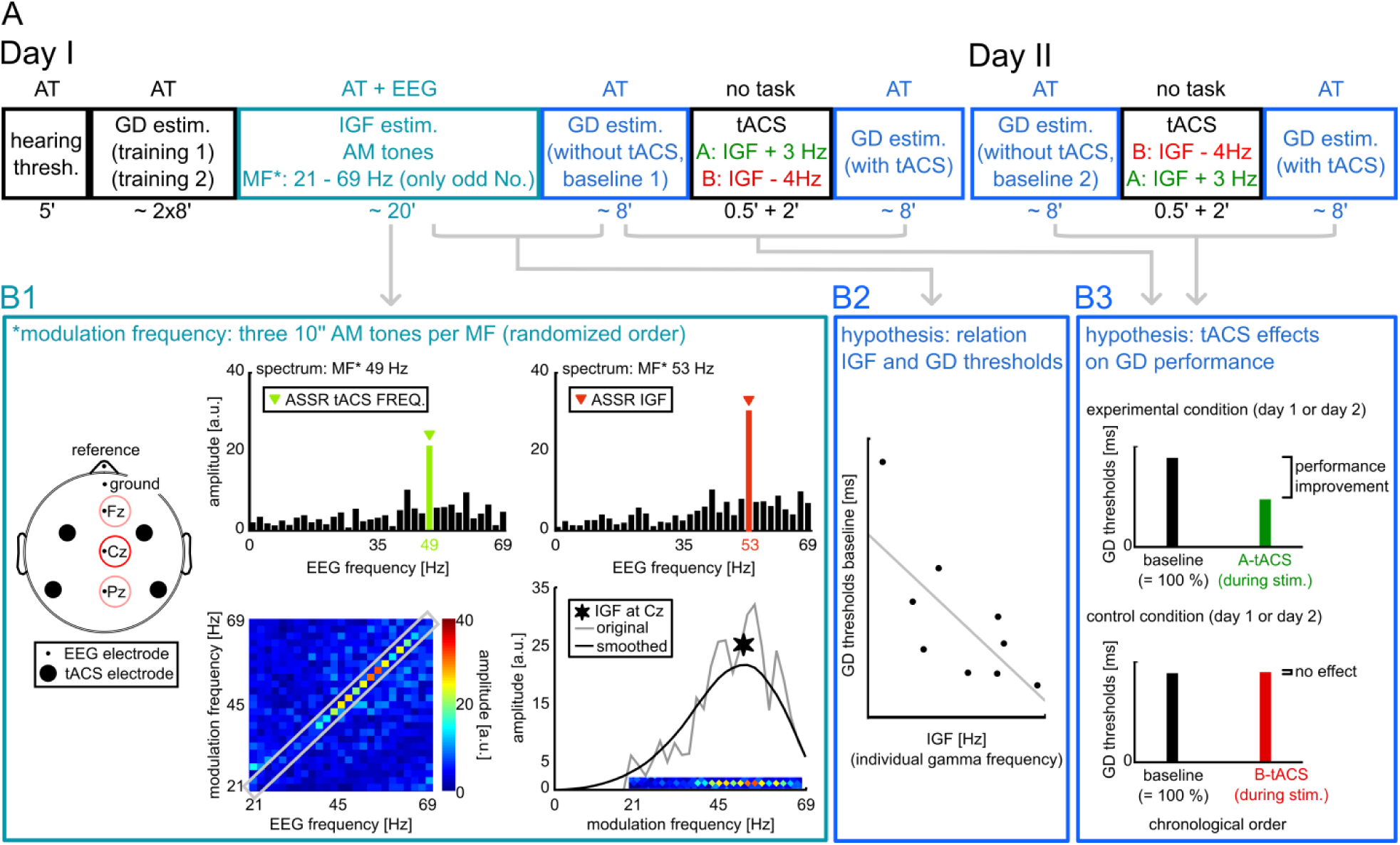
Timeline, experimental setup and hypotheses. A) Timeline of the experiment over two days. Type of task is specified above the corresponding box (AT: auditory task, AT + EEG: auditory task during electroencephalographic (EEG) recording). Duration of single parts are given below the timeline in minutes (‘). After a rough measurement of individuals hearing thresholds (hearing thresh.) each participant underwent two trainings of a between-channel gap detection (GD) estimation task followed by an auditory task for which amplitude modulated (AM) tones with modulation frequencies from 21 to 69 Hz (in steps of 2 Hz) were presented. Following these preparatory parts, participants performed the same GD task as estimation for baseline 1 (without tACS) and, after 2.5 minutes of transcranial alternating current stimulation (tACS) without any behavioral task, GD performance was reassessed during tACS. At day two, participants again performed the GD task for a second baseline estimation and underwent the same tACS procedure as at day one (2.5 minutes without task and then with GD task). TACS frequency for the experimental condition was IGF + 3 Hz and for the control condition it was IGF - 4 Hz. Conditions were randomized across participants. B) Grey arrows indicate which box belongs to which part of the experiment. In box 1 (B1), a rough scheme of the IGF estimation is given. AM tones were 10 seconds (‘’) long and for each modulation frequency three AM tones were presented (in a randomized order). EEG electrodes Fz, Cz and Pz were recorded during AM tone presentation (positions of tACS electrodes are depicted for information only) and data analysis was conducted for each electrode. Exemplary data set of electrode Cz: spectra of recorded EEG data are shown in the upper row of B1: clear spectral peaks (marked by triangles) depict auditory steady state responses (ASSRs) to modulation frequencies of the AM tones (49 and 53 Hz). In the lower row, spectra of all modulation frequencies are combined in one square with ASSR amplitudes being color coded (left). The extracted diagonal depicts amplitudes of all ASSRs (right). The maxima of the smoothed diagonals were compared across electrodes and the frequency of maximum of the maxima was taken as individual gamma frequency as indicated by a star. In box 2 (B2), the hypothesized negative correlation between IGF and GD performance at baseline (higher IGFs are related to better GD performance) is depicted. In box 3 (B3), hypothesized effects of tACS on GD performance are depicted: in the experimental condition, a performance improvement for GD performance during tACS is expected whereas in the control condition no effect of tACS on GD performance is expected.

Participants performed six sessions of a GD task. Session one and two were training sessions to familiarize the elderly participants with the task as well as to reduce learning effects which are known to influence the estimated GD thresholds (Smith *et al*., 2008). If necessary, verbal instructions were given additionally to the information sheet during the first session of the GD task. Session three was used as baseline one (BL1) at day one and served as 100% marker for session four during which participants were electrically stimulated. At day two, session five served as baseline two (BL2) and 100% marker for the session six in which participants were again electrically stimulated. GD thresholds of BL1 and BL2 were correlated to test for test-retest reliability with Spearman’s rank correlation coefficient.

Auditory stimuli were presented via a digital-analog-converter (NI USB 6229, National Instruments Germany GmbH, Munich, Germany), a Programmable Attenuator PA5 and the Headphone Driver HB7 (both Tucker Davis Technologies, Alachua, USA). Binaural air-pressure headphones (E-A-RTONE Gold 3A Insert Earphones, 3 M Auditory Systems, Indianapolis, USA) were used to keep electric devices as far away from the EEG electrodes as possible. Presentation of auditory stimuli and visual information was controlled with MATLAB 2012a (64bit, The Mathworks, Natick, USA), the psychophysics toolbox (3.0.11) and the daq toolbox (both implemented in MATLAB).

Recorded behavioral data were analyzed in Matlab 2012a (The Mathworks, Natick, USA). EEG data analysis was performed using EEGLAB (Delorme & Makeig, 2004).

### Electrical stimulation

In the domain of transcranial electric current stimulation, recent methodological developments allow modelling of the current flow through the head (Wagner *et al*., 2014) during stimulation. In Baltus *et al*. (2017) an optimized stimulation electrode setup specifically suitable for auditory tACS was introduced (Fig. 3). With this setup we demonstrated a frequency-specific effect of auditory tACS on auditory temporal resolution in normal hearing adults. This electrode setup (EEG positions FC5 and TP7/P7 for the left-hemisphere channel and FC4 and TP8/P8 for the right-hemisphere channel) was computed using a highly realistic finite element methods modelling approach based on the findings of (Wagner *et al*., 2016b). For the modelling, targets in the auditory cortices were determined by estimating dipoles of auditory N1 measured in a combined auditory evoked potential/auditory evoked field study in both auditory cortices for one exemplary subject (not part of the study cohort). In addition, a highly realistic geometry-adapted hexahedral FEM mesh of the head of the exemplary subjects was created based on a high resolution multimodal MRI (T1-weighted, T2-weighted, DTI). A mathematical problem was formulated with the aim to find an optimal stimulation pattern producing maximal current densities in target areas (while they are minimal in the remaining areas of the brain) and current flow that is parallel to the defined targets. The optimal stimulation pattern (allowing 74 possible electrode locations based on an extended 10–10 EEG electrode system) was calculated, by finding a unique solution for the mathematical problem. After finding the optimized stimulation pattern (one specific value for the 74 electrode positions, Fig. 3A), the current densities and current flow directions were recalculated using two stimulation electrodes, one anode and one cathode, per hemisphere (Fig. 3B). Recalculation showed desired current density patterns in auditory cortex areas (Fig. 3C).

**Figure 3:**
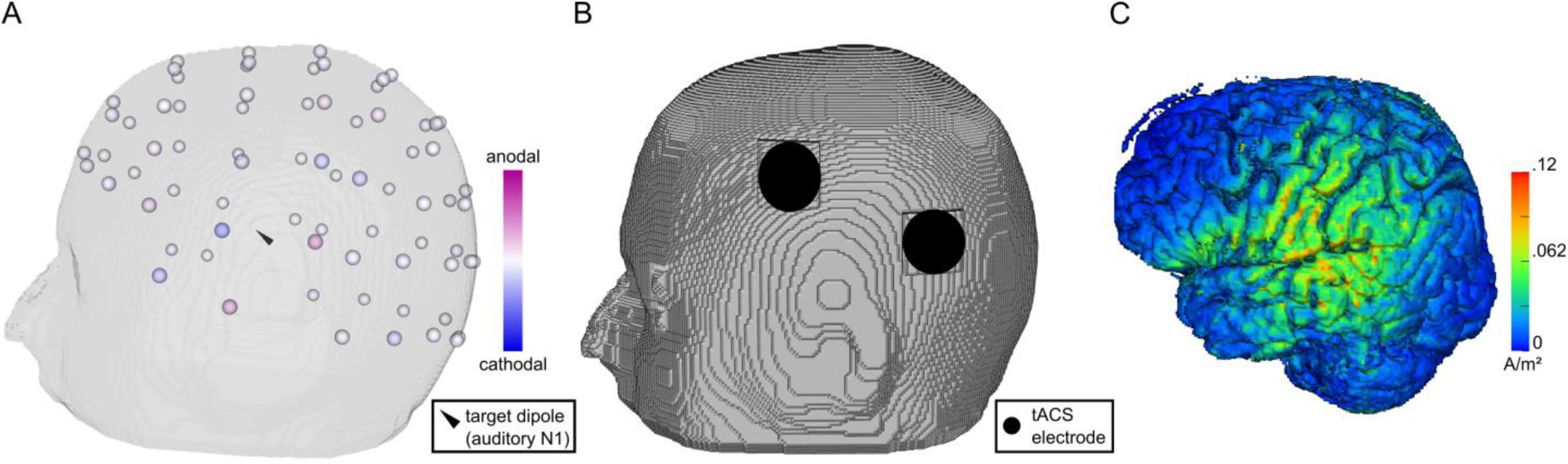
Auditory transcranial alternating current stimulation (tACS) electrode setup and results of current flow modelling for the left hemisphere (right hemisphere: analogous). A) Target dipoles (estimated from auditory N1 measured in a combined auditory evoked potential/auditory evoked field study) in auditory areas (black triangles) are used to compute an optimal tACS stimulation pattern allowing 74 stimulation electrodes (extended 10–10 EEG system). Each electrode can be either anodal or cathodal with anodic (cathodic) currents summing up to +/− 2 mA (for details see Baltus *et al*., 2017 and Wagner, Burger, *et al*., 2016). B) Simplified stimulation electrode setup (only two electrodes per hemisphere). C) Finite element modelling of current densities in the left hemisphere for the chosen two-electrode setup demonstrates considerable current densities in auditory cortex and adjacent areas.

Electrical stimulation was applied during session four and session six of the GD task (Fig 2A). A neuroConn multichannel stimulator (DC-STIMULATOR MC, neurConn GmbH, Ilmenau, Germany) with two channels, one for each hemisphere, was used. For each channel, two round electrodes with 2.5 cm diameter were attached using alcohol for pre-attachment cleaning, a mix of ABRALYT gel (Abrasive electrolyte-Gel, EASYCAP, Herrsching, Germany), and Ten20 Conductive paste (Weaver and company, Aurora, USA). With this procedure, impedances for both channels were kept below 10 kQ for each participant. The current was ramped in and out to avoid sensation of current flow. Stimulation strength of 1 mA (peak to peak) was reached after 30 seconds. During a waiting period of two minutes, after the final stimulation strength was reached, participants were instructed to sit relaxed. The stimulation was immediately switched off after the GD task had ended (Fig. 2A). On average the stimulation duration was 8.5 ± 1.8 minutes.

### Individual gamma frequency (IGF) estimation

Amplitude modulated tones with a center frequency of 1000 Hz and modulation frequencies from 21 Hz to 70 Hz (in steps of two Hz) were presented three times for ten seconds in a randomized order (Fig. 2B1). In a previous study, modulation frequencies within the same range (21 Hz to 70 Hz) but with steps of one Hz were used to estimate the IGF (Baltus *et al*., 2017). For that dataset, test-retest reliability for IGF estimated with modulation frequencies in steps of one Hz and steps of two Hz were found to be sufficiently high *(Spearman’s rho* = 0.92, *p* = 2e-11). Hence, decreasing the duration of data recording for elderly people, by using modulation frequencies in steps of two Hz, is justified. Starting phase angles of the amplitude modulated tones were zero degree to avoid spectral splatter at the tone onset. Analogously to the GD stimuli, the sampling frequency of the ASSR stimuli corresponded to 10 times the EEG sampling frequency. The modulation depth was 100 % and ASSR stimuli were presented 60 dB above individual hearing threshold which has been shown to elicit reliable ASSRs around the preferred frequency (Baltus & Herrmann, 2015). An auditory vigilance task was used to ensure a stable level of attention (Ross *et al*., 2004): one third of the trials were characterized by a loudness decrease of 30 % of the normal stimuli and served as target trials that had to be indicated by a button press of the subjects. The loudness decrease occurred between second 4 and 8 and lasted for one second. Participants received feedback instantly after each button press in order to keep their motivation on a constant level (Deci *et al*., 1999). Eye-blink artifacts were avoided by instructing the subjects to close their eyes during each stimulus presentation.

The individual tACS frequency was determined based on the individual EEG data of each modulation frequency (3x10 seconds) and each electrode. The EEG data were epoched in 1-second segments with an overlap of 500 ms based on triggers marking the troughs of the amplitude modulation. The first two epochs were not considered for analysis to ensure a total cut-off of the auditory onset response to the stimuli (Röschke *et al*., 1996). Remaining epochs were baseline corrected with respect to the whole epoch and averaged to calculate an event related potential (ERP) for each modulation frequency (Fig. 2B1). Spectra of each ERP were calculated with the Fast Fourier Transform and 1/f-corrected by multiplying each spectral value with its frequency to control for the 1/f-characteristic of EEG spectra (Herrmann, 2001). Amplitudes of the auditory steady state responses (ASSRs; John & Picton, 2000; Picton *et al*., 2003) were extracted for each modulation frequency and resulted in an individual ASSR profile for each electrode (Fig. 2B1). ASSR profiles were smoothed with a simple moving average filter implemented with the MATLAB function filtfilt.m to avoid phase distortions. In a previous study, we discovered a possible dependence of ASSR profile on the orientation of the conjectured sources of the ASSR in the auditory cortex (Baltus & Herrmann, 2015). Therefore, the parameters of the maxima of the ASSR profile (frequency and amplitude) were compared between all electrodes (Fz, Cz, and Pz). The modulation frequency that elicited the largest ASSR amplitude in one of these three electrodes was used as estimation for the IGF. TACS frequencies where then chosen to be close to the IGF.

Determination of the tACS frequencies was based on three constraints: first, the concept of the Arnold tongue, which suggests choosing a driving frequency close to the endogenous frequency if weak external driving forces (such as tACS) are applied (Pikovsky *et al*., 2001; Notbohm *et al*., 2016). Second, the explanatory model of Vossen (2015) for the observed phenomenon that tACS with frequencies slightly *below* the endogenous frequency do not lead to entrainment, i.e. an increase in amplitude at the driving frequency, but rather to an increase in amplitude at the endogenous frequency. The model is supported by electrophysiological and behavioral findings in Baltus *et al*. (2017) where auditory tACS 4 Hz *below* IGF had no effect on either GD performance or post-tACS ASSR amplitudes. Third, the entrainment region hypothesis (McAuley *et al*., 2006) which hypothesizes a decrease in width of entrainment regions with age, meaning that Arnold tongues become smaller with age. Therefore, higher driving forces (i.e., stronger intensities), which are unfeasible with tACS, or driving frequencies closer to the endogenous frequency are required.

Based on the three constraints introduced in the introduction, the tACS frequency for the experimental condition was chosen to be *3 Hz above IGF* to satisfy the constraints of the entrainment region hypothesis and the Arnold tongue (in a recent study, young adults showed increased GD performance during tACS with stimulation frequencies *4 Hz above IGF* (Baltus *et al*., 2017).

The tACS frequency of the control condition was chosen to be *4 Hz below IGF* of the participants. TACS at *4 Hz below IGF* is expected to be ineffective with regard to GD performance (based on the explanatory model of Vossen (2015), the behavioral and electrophysiological results of Baltus *et al*. (2017) and the idea that driving frequencies further apart from the endogenous frequency are less effective to modulate ongoing oscillations (Pikovsky *et al*., 2001).

### GD threshold estimation

Stimuli for the estimation of the between-channel gap detection thresholds were created based on the methods introduced by Phillips (1997) and their modifications by Baltus & Herrmann (2015). Band-passed filtered (48 dB/oct) white noises with two different center frequencies (2000 Hz and 1000 Hz) were used as leading (sound before the gap) and trailing sound (sound following the gap), respectively. Leading and trailing noises were essentially the same white noise vector filtered with two different band-pass filters (0.125 octave below and above center frequency) and normalized to the maximum of each filtered noise. Each noise was multiplied by a Tukey window with a half-cosine shaped rise- and fall-time of 2 ms to avoid spectral splatter (Plack, 2013). The sampling frequency of the stimuli was 50,000 Hz, i.e. tenfold the EEG sampling frequency (5000 Hz). Total length of one stimulus was one second. Gap length was divided in half and subtracted from leading and trailing noise to ensure equal length of both noises. Gap lengths ranged from 1 ms to 250 ms in steps of 0.1 ms. GD stimuli were presented relative to the individual hearing threshold 40 dB above the absolute threshold of hearing of each participant.

Thresholds for GD performance of each subject were estimated by using a two-alternative forced choice task in an adaptive staircase procedure with an a-priori probability of 0.5 for the gap appearing in the first or the second stimulus. The used staircase procedure was introduced by Garcia-Perez (García-Pérez, 1998) and is especially suitable for the small sample staircase applied in this experiment with only 14 reversal points. The maximal gap size and starting value for the staircase procedure was 250 ms which ensures a clear perception of the gap. The arithmetic mean of the gap sizes at the last 12 reversals was used as the GD threshold estimation.

### Statistical analyses

Based on the assumption that higher endogenous frequencies in auditory cortex (higher IGFs) lead to better GD performance (resulting in a negative correlation between the two variables), GD thresholds estimated in the two baseline sessions were individually correlated with the IGFs using Spearman’s rho with the alternative hypothesis of a correlation less than zero (left-tailed, Fig. 2B2).

Effects of tACS on gap detection were calculated by computing the percentage of change from session three (baseline 1) to session four (stimulation session) as well as the percentage of change from session five (baseline) to session six (stimulation session; Fig. 2B3) for each stimulation condition, respectively. Relative changes between baseline GD performance and GD performance during tACS were calculated with respect to baseline GD performance being. Subsequently, 100% was subtracted from each resulting percentage value (resulting in a 0% value, if there was no change from baseline GD estimation to GD estimation during tACS). In both conditions, relative changes were tested for outliers (the two most extreme values) with Dixon’s Q-test (Dixon, 1951). Relative changes were then tested for normality with the Kolmogorov-Smirnov test and compared between stimulation conditions (Fig. 2B3) using the Wilcoxon rank sum test with an alpha level of 0.05.

To discover a possible relationship between the effect of tACS on auditory temporal resolution (i.e., gap detection) and the endogenous frequency of the auditory cortex of the elderly individuals, the baseline corrected individual GD thresholds (relative changes) were correlated with the IGF using the non-parametric Spearman’s rho (with an undirected alternative hypothesis). For the experimental condition, a non-zero, significant correlation coefficient was expected. For the control condition (no effect of tACS expected), a zero correlation was expected.

## Results

Average pure tone audiograms demonstrate considerable hearing abilities above 25 dB for all subjects (Fig. 1). In addition, average pure tone audiograms correlated significantly (*Spearman’s rho* = −0.78, *p* = 0.004) with the average hearing thresholds estimated with the adapted version of the self-adjustable adaptation procedure from Carhart and Jerger (Carhart & Jerger, 1959). This indicates a link between good performance in pure tone audiograms and good performance in hearing threshold estimation procedures using white noise as stimulus. Therefore, our adapted hearing threshold estimation is a valid procedure to correct for individual differences in hearing abilities and auditory stimuli were perceived at a similar loudness level.

GD thresholds estimated as baseline for the GD performance during tACS correlated significantly with the IGFs: *Spearman’s rho* = −0.67, *p* = 0.03 for the correlation between IGF and baseline 1 measured in GD session three and *Spearman’s rho* = −0.67, *p* = 0.03 for the correlation between IGF and baseline 2 measured in GD session five (Fig. 4). The similarity between the two correlations is attributable to the high test-retest reliability of GD thresholds estimated as baseline 1 and as baseline 2 (Fig. 4, *Spearman’s rho* = 0.91, *p* < 0.001). These two observations (the high test-retest reliability of GD thresholds at baseline and the significant negative correlation between IGF and GD performance at baseline, Fig. 3) nicely replicate the correlations found in Baltus and Herrmann (2015).

**Figure 4:**
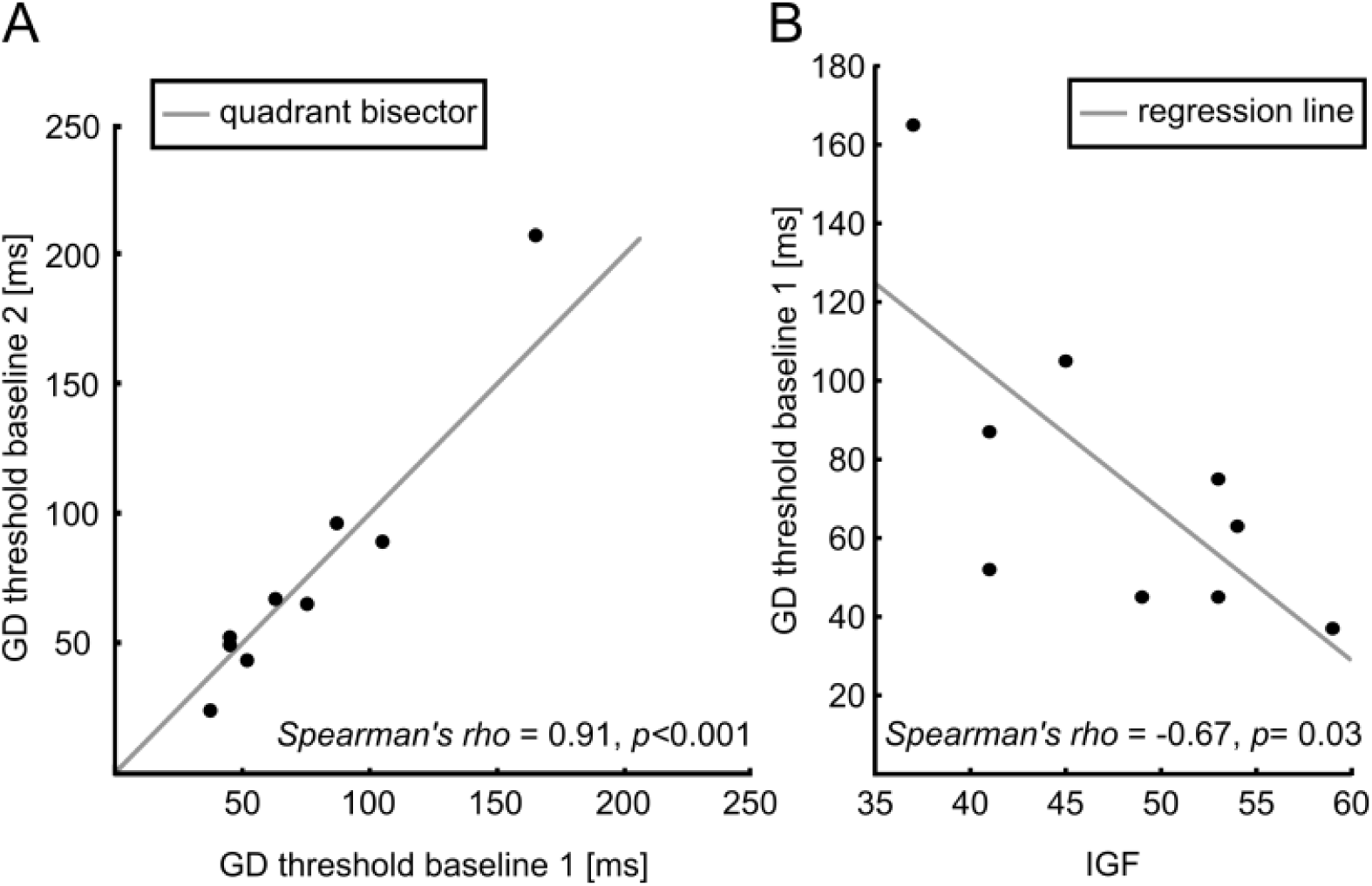
Test-retest reliability of gap detection (GD) thresholds and correlation between individual gamma frequencies (IGFs) and baseline gap detection (GD) thresholds. A) High test-retest reliability of GD threshold measured at day one of the experiment as baseline for the first stimulation session and at day two as baseline for the second stimulation session can be identified as sufficiently high. Grey line indicates the quadrant bisector (line through the origin with slope one). B) As hypothesized, IGFs correlated significantly with GD thresholds at baseline. For illustrational reasons, IGFs are only correlated with GD thresholds measured as baseline for the first stimulation session. However, statistical values for the correlation between IGF and GD thresholds measured as baseline for the second stimulation session are exactly the same due to high test-retest reliability of baseline GD thresholds.

In contrast to the hypothesis derived from the results of younger adults, where a significant difference between stimulation groups were found (Baltus *et al*., 2017), relative changes between conditions in the current study were found not to be significantly different. However, for the experimental condition (tACS frequency = IGF + 3 Hz), we found a significant positive correlation between IGF and change in performance (*Spearman’s rho* = 0.79, *p* = 0.01, Fig. 5A). For the control condition (tACS frequency = IGF - 4 Hz), however, no correlation was observed between IGF and change in performance (*Spearman’s rho* = 0.02, *p* = 0.9, Fig. 5B). In neither condition, outliers could be identified with Dixon’s Q-test (experimental condition: r_11_ = 0.0627<0.634, α = 0.01; control condition: r_11_ = 0.5186<0.634, α=0.01).

**Figure 5:**
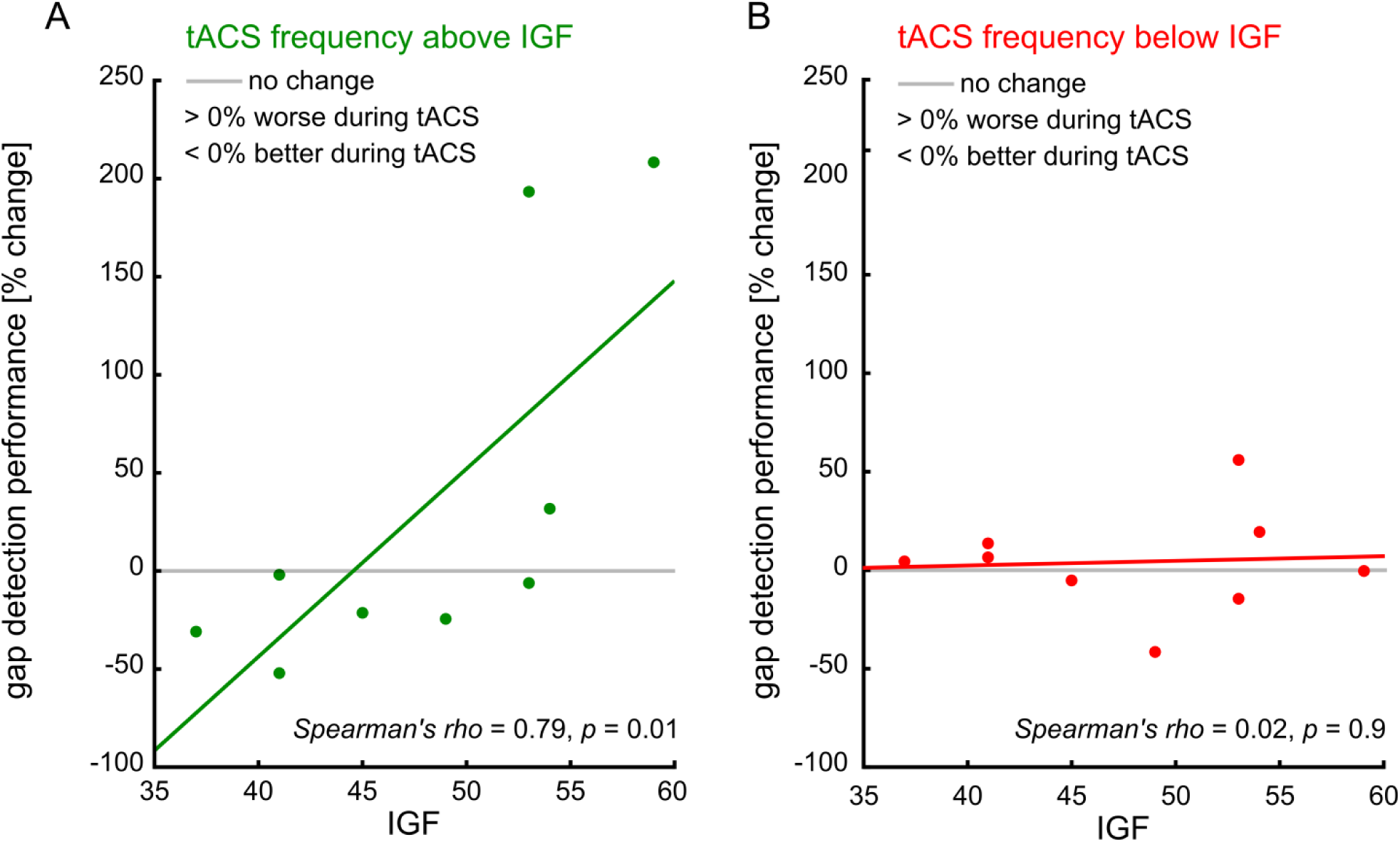
Results of transcranial alternating current stimulation (tACS) effects. A) In the experimental condition, the effect of tACS is correlated with individual gamma frequencies of the participants: Participants with lower IGFs profit from tACS (performance increases in comparison to baseline, percentage value below zero) whereas participants with higher IGFs are hampered by tACS (performance decreases in comparison to baseline, percentage value above zero). B) In the control condition (no effect of tACS was hypothesized), participants did not change performance during tACS in comparison to baseline.

## Discussion

In the current study, the role of oscillatory activity in auditory cortex for auditory processing with regard to auditory temporal resolution was investigated and further established. Recent studies on GD processing led to the idea of a potential accelerating effect of auditory tACS for elderly people for which a general decline in auditory processing speed had been observed. We hypothesized an enhancement of GD performance during auditory tACS with stimulation frequencies above IGF in the experimental condition. What we found was an interaction between IGF and GD performance during auditory tACS. The experimental condition was contrasted with a control condition in which auditory tACS with stimulation frequencies below IGF was expected to have no effect on GD performance. The results of the current study supported the hypothesis concerning the control condition: no effect of auditory tACS on GD performance was observed in the control condition. All subjects participated in both conditions to address the problem of large variability of sensory processing performance observed in elderly people. In addition, test-retest reliability of GD thresholds was assessed to control for variability in performance which could be an alternative explanation for the observed results. We found test-retest reliability of GD thresholds to be sufficiently high in order to consider them as an estimate for a physiological trait. We conducted rough, frequency unspecific hearing thresholds at the beginning of the experiment for each participant. In addition, participants were pre-screened by the Hörzentrum Oldenburg with a pure tone audiometry. We found the roughly estimated hearing thresholds to correlate significantly with averages of pure tone audiograms indicating that the method used in the current study is a reasonable proxy for the more complex, lengthy pure tone audiometry.

Results of the current study reveal a more complex interaction between IGF and the effect of auditory tACS on individual gap detection thresholds for older adults than for younger adults. In contrast to younger adults, in older adults effects of auditory tACS seem to depend on frequencies of ongoing gamma oscillations: participants with lower IGF profit from auditory tACS in the experimental condition (tACS frequency: 3 Hz above IGF), whereas participants with higher IGFs where hampered by tACS in the experimental condition. No effect of tACS on GD performance was observed in the control condition (tACS frequency: 4 Hz below IGF).

Since GD performance at baseline showed high test-retest reliability and, in addition, the tACS effect was only present in the experimental condition (although subjects were stimulated in both conditions) particular change of GD performance during the application of auditory tACS in the experimental condition can be considered a consequence of the applied stimulation. This observation is in line with results of other studies in which a change in behavioral performance was attributed to a modulation of ongoing neuronal oscillations in relevant brain areas due to the application of tACS (Neuling *et al*., 2012a; Samaha & Postle, 2015; Rufener *et al*., 2016a; Rufener *et al*., 2016b). The basic aim of the application of tACS in these studies is to prove that brain oscillations not only represent incidentally regular voltage changes at the surface of the brain but are directly connected to processing mechanisms in the brain and therefore physiologically relevant. Application of tACS is supposed to influence endogenous brain oscillations in order to change the frequency of these oscillations (Fröhlich, 2015). A subsequent change in behavioral performance can therefore be interpreted as a consequence of the frequency change. Auditory tACS has successfully been applied to modulate specific processing characteristics leading to altered behavioral performances in young adults (Riecke *et al*., 2015; Rufener, Zaehle, *et al*., 2016; Baltus *et al*., 2017) and older adults (Rufener *et al*., 2016a). The rationale of a direct link between brain oscillations and physiologically relevant processing mechanisms is still based on indirect observations but these observations more strongly support a causal relationship between brain oscillations and behavior than purely correlational observations (Herrmann *et al*., 2016).

The rationale for the chosen tACS frequencies in the experimental and the control condition is supported by human and animal modelling studies. TACS *below* the frequency of endogenous oscillations leads to an increase in amplitude *at* the frequency of endogenous oscillations whereas tACS *above* the frequency of endogenous oscillations leads to a frequency *shift* of the endogenous oscillation towards the applied frequency. This argumentation is based on human data (Vossen *et al*., 2015) showing that tACS below the endogenous frequency led to an amplitude increase in post-stimulation EEG measurements at the endogenous frequency, as well as in *in-vitro* data showing that the frequency of light-controlled oscillatory activity in mouse neocortical slices was only possible to be shifted towards slightly higher frequencies (Schmidt *et al*., 2014).

We found a significant relation between GD performance and IGF where participants with lower IGFs benefited from accelerating tACS whereas participants with high IGFs were hampered by the accelerating tACS. Participants at the upper end of the IGF scale ranked among the best performing subjects which is consistent with the idea that faster endogenous oscillations are the physiological basis for increased auditory temporal resolution.

It has been observed in other transcranial stimulation studies that perceptual processing deteriorates if associated oscillatory activity is modulated, especially if the linked neuronal process is already initially at a very high level (Schwarzkopf *et al*., 2011; Neuling *et al*., 2013; Heimrath *et al*., 2014). Since IGFs tend to decrease with age (Purcell *et al*., 2004), the participants who showed decreased performance during tACS in our study appeared to have IGFs at the upper bound of the scale. Thus, the auditory system of those subjects might perform already at an absolute upper limit of related processing mechanisms where any influence can only lead to a degradation of processing success. Such a deteriorating effect of gamma tACS (40 Hz) on processing of rapidly acoustic features was also observed by Rufener *et al*. (2016b). In line with our observation of a trait dependent effect of brain stimulation, a study, investigating the effects of brain stimulation on working memory performance in elderly found the direction of the effect (improvement vs. impairment) to be dependent on participant’s educational levels (Berryhill & Jones, 2012). However, due to lower sample size in the current study and only moderate effect sizes of 0.33 in a previous study, where effects of different tACS frequencies on gap detection thresholds in younger adults were investigated (Baltus *et al*., 2017), observed effects might not be generic and further research is necessary to disentangle frequency specific effects of tACS on gap detection in healthy elder adults. In addition, all subjects demonstrate especially good hearing performance in daily life (see average PTA, fig. 1) and therefore, the cohort is probably less representative than a cohort, that would include elderly people with hearing loss.

Human (Crone *et al*., 2001; Schadow *et al*., 2007; Lenz *et al*., 2008; Herrmann *et al*., 2010) as well as in in-vivo and in-vitro studies in animals (Demiralp *et al*., 1996; Brosch *et al*., 2002) revealed auditory stimulus-evoked and -induced gamma activity after stimulus onset. In humans, peak frequencies of stimulus-evoked sensory gamma activity are linked to the IGF as estimated in the current study (Zaehle *et al*., 2010). In a different study, in which GABA concentrations at baseline in sensory areas were correlated with peak frequencies of stimulus-induced gamma band responses, results revealed a positive relation between GABA and peak frequency, with higher GABA concentration being linked to higher gamma peak frequencies (Muthukumaraswamy *et al*., 2009). The relation between GABA and frequency of network frequency is also indicated by a model using repetitively firing interneurons and dense inhibitory interconnectivity (Traub *et al*., 1996; Whittington *et al*., 2000). Recently, due to the large quantity of contrary findings in tACS studies, a dependency of tACS effects on the balance between excitatory and inhibitory brain activity at baseline has been hypothesized (Krause & Kadosh, 2014). Under the provision outlined in this paragraph, our tACS findings, with tACS effects depending upon frequency of gamma activity at baseline, support this idea.

The tACS setup used in this study is based on finite element modelling simulations suggesting sufficient current flow during tACS in auditory cortex areas (Wagner, Lucka, *et al*., 2016; Baltus *et al*., 2017). Based on the results of the stimulation and the behavioral observations in the current study, a modulation of relevant (oscillatory) activity due to tACS is likely to be the underlying physiological mechanism for the observed change in GD performance. Hence, the possibility to manipulate GD performance with tACS can be interpreted as evidence for a physiological relevance of neuronal oscillations in the gamma range for auditory temporal resolution expressed by GD thresholds. Consequently, this verifies on a more corroborated level than pure correlational observations the relevance of neuronal oscillations for sensory processing.

In summary, results of the current study further support the evidence for a causal relationship between auditory temporal resolution expressed by GD thresholds and neuronal oscillation in auditory cortex areas. In general, this is the first study using tACS with an individualized stimulation protocol to investigate in a causal way the relation between oscillatory gamma activity in the auditory cortex and temporal resolution of the cortical auditory system in elderly people. Two important results can be concluded: first, individual auditory temporal resolution is directly related to the frequency of neuronal oscillations in sensory processing areas and second, if successful modulation of sensory performance is aspired with tACS, individualized tACS protocols are essential.

## Acknowledgements

This study was supported by the German Research Foundation (DFG grant SFB/TRR 31). We want to thank Julia Schmidt for her help with data acquisition.

## Authors contributions

Conceived and designed the experiments: AB, CSH. Performed the experiments: AB. Analyzed the data: AB, CSH. Wrote the paper: AB, CSH.

## Abbreviations

ASSR: auditory steady state response
EEG: electroencephalography
ERP: event related potential
GABA: gamma-Aminobutyric acid
GD: gap detection
Hz: Hertz
IGF: individual gamma frequency
mA: milliampere
MF: modulation frequency
ms: milliseconds
PTA: pure tone audiometry
tACS: transcranial alternating current stimulation

